# Origins and long-term patterns of copy-number variation in rhesus macaques

**DOI:** 10.1101/749416

**Authors:** Gregg W.C Thomas, Richard J. Wang, Jelena Nguyen, R. Alan Harris, Muthuswamy Raveendran, Jeffrey Rogers, Matthew W. Hahn

## Abstract

Mutations play a key role in the development of disease in an individual and the evolution of traits within species. Recent work in humans and other primates has clarified the origins and patterns of single nucleotide variants, showing that most arise in the father’s germline during spermatogenesis. It remains unknown whether larger mutations, such as deletions and duplications of hundreds or thousands of nucleotides, follow similar patterns. Such mutations lead to copy-number variation (CNV) within and between species, and can have profound effects by deleting or duplicating genes. Here we analyze patterns of CNV mutations in 32 rhesus macaque individuals from 14 parent-offspring trios. We find the rate of CNV mutations per generation is low (less than one per genome) and we observe no correlation between parental age and the number of CNVs that are passed on to offspring. We also examine segregating CNVs within the rhesus macaque sample and compare them to a similar dataset from humans, finding that both species have far more segregating deletions than duplications. We contrast this with long-term patterns of gene copy-number evolution between 17 mammals, where the proportion of deletions that become fixed along the macaque lineage is much smaller than the proportion of segregating deletions. These results suggest purifying selection acting on deletions, such that the majority of them are removed from the population over time. Rhesus macaques are an important biomedical model organism, so these results will aid in our understanding of this species and the disease models it supports.

## Introduction

Mutations are the source of all genetic variation, and can have both immediate effects on individual phenotypes and lasting impacts on genome evolution. Understanding how mutations arise and spread through a population in the short-term can therefore aid our understanding of disease, while understanding their effects in the long-term aid our understanding of evolution in populations and species. Recent work in humans and other primates have unveiled patterns of mutation for single nucleotide variants using pedigrees of related individuals. For instance, studies in primates have found a strong paternal age effect on the number of *de novo* single nucleotide mutations: older fathers tend to pass on more mutations (Kong et al. 2012; Venn et al. 2014; Jonsson et al. 2017; Thomas et al. 2018). This is likely due to a combination of errors accruing from both ongoing spermatogenesis and unrepaired DNA damage. However, no such paternal age effect has been found among *de novo* deletions and duplications (also known as copy-number variants, or CNVs) in humans (MacArthur et al. 2014; Kloosterman et al. 2015; Brandler et al. 2016; Girard et al. 2016), though the origin of CNVs have been studied less often than single nucleotide mutations (Sebat et al. 2007; Itsara et al. 2010; Schrider et al. 2013; MacArthur et al. 2014; Kloosterman et al. 2015; Brandler et al. 2016; Girard et al. 2016; Werling et al. 2018).

The frequency of CNVs among lineages and the density of CNVs along the genome have been found to be highly variable among primates (Fortna et al. 2004; Jiang et al. 2007; Sudmant et al. 2010; Gazave et al. 2011), with CNV hotspots in multiple species having been described (Perry et al. 2006; Perry et al. 2008; Gokcumen et al. 2011). Duplications in genic regions have been found to outnumber deletions in many lineages when comparing closely related species (Fortna et al. 2004; Dumas et al. 2007; Sudmant et al. 2013), possibly indicating differential effects of natural selection on gene duplications versus deletions. However, recent whole-genome studies in humans point to a different pattern in non-genic regions, with deletions far outnumbering duplications (Brandler et al. 2016). In order to determine whether such patterns are specific to humans, or are representative of the joint effects of mutational input and selection on the long-term survival of duplications and deletions, we require fine-scale studies in additional species.

Rhesus macaques (*Macaca mulatta*) are a widely used model organism, especially for studies of human diseases. Understanding the underpinnings of genetic variation in this species may help to enhance studies on existing or new disease models, in addition to aiding our understanding of the genetic basis of evolutionary change. Previous studies of rhesus macaque CNVs have used array-based comparative genomic hybridization (aCGH) to detect events and have found that the frequency of duplications either matches or exceeds that of deletions (Lee et al. 2008; Gokcumen et al. 2011). However, aCGH methods are limited in their detection of short deletions and duplications (Medvedev et al. 2009; Zarrei et al. 2015). Patterns of variation in CNVs shorter than the detectable limit by aCGH remain uncharacterized. Read-based methods—which use read depth, read orientation, discordance of paired-end reads from a reference genome, or a combination of these signals (reviewed in Medvedev et al. 2009; Zhang et al. 2019)—will help to clarify patterns of duplication and loss.

Here, we use deep sequencing of 32 rhesus macaques including 14 sire-dam-offspring trios to uncover patterns of copy-number mutation and variation in this species. We find that, contrary to aCGH studies, deletions make up the vast majority of polymorphic CNVs within rhesus macaques. Using unrelated individuals, we find that patterns of segregating CNVs are similar between macaques and humans. By sequencing parent-offspring trios we are also able to investigate the occurrence of *de novo* CNVs. We find that the number of *de novo* CNVs per generation is less than one per genome in both macaques and humans, and that parental age has no detectable effect on the rate of these types of mutations in either species. Finally, we compare patterns of deletions and duplications in our sample to those of long-term gene gains and losses along the lineage leading to macaques from their last common ancestor with baboons (genus *Papio*). Interestingly, while deletions make up the vast majority of polymorphisms in our sample, the number of genes gained and lost along the macaque lineage are roughly equal. These patterns give us a first look at the origins of copy-number variation using whole genome sequencing in a nonhuman primate, and will help improve modeling of these types of mutations in relation to both disease prediction and evolutionary analyses.

## Results

### Patterns of copy-number variation in rhesus macaques

We identified CNVs by sequencing the whole genomes of 32 Indian-origin rhesus macaques (Figure 1A; Wang et al. 2019). We mapped the reads from these samples to the reference macaque genome (rheMac8, also called Mmul_8.0.1, downloaded April 12, 2018) and identified CNVs based on split and discordant read patterns using Lumpy (Layer et al. 2014), SVtyper (Chiang et al. 2015), and SVtools (Larson et al. 2018). We then filtered these calls by read-depth using Duphold (Pedersen and Quinlan 2019). In total we found 3,313 deletions and 441 duplications among these 32 individuals relative to the reference genome, meaning that roughly 88% of variants segregating in our sample are deletions (Figure 1B). This is in stark contrast to previous CNV studies in rhesus macaques, which found roughly half of events to be deletions and half to be duplications (Gokcumen et al. 2011). One possibility for this difference is that the previous study could not resolve events shorter than a few kilobases (minimum length 3518 bases), while the read-based methods employed here can. This contrast from an increased level of resolution is consistent with studies in *Drosophila melanogaster* that found a bias toward deletions only for short events (Schrider et al. 2013).

**Figure 1:**
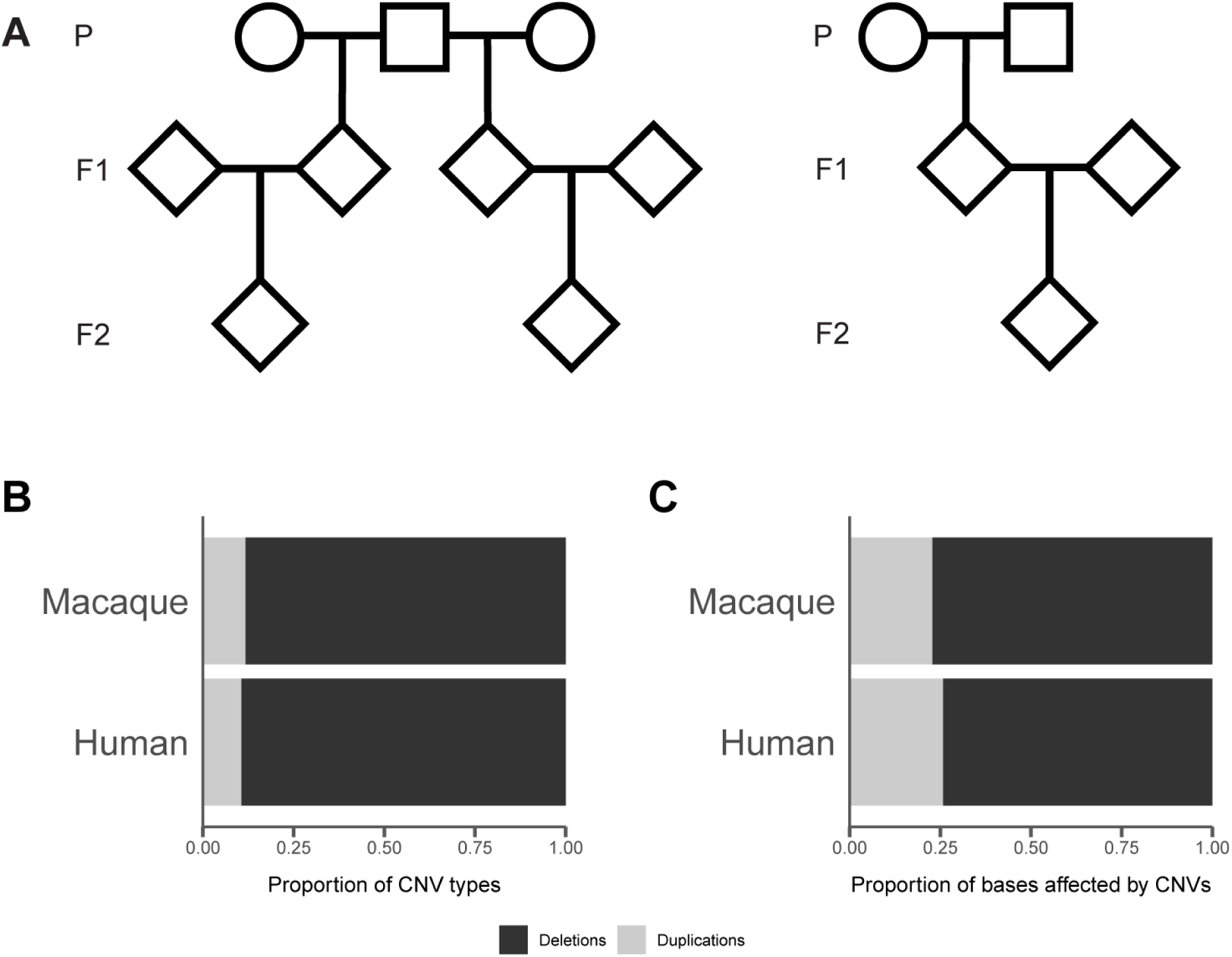
(**A**) Pedigrees of sequenced macaques. The 14 trios were contained within 3 families similar to the one on the left, and 1 family similar to the one on the right. (**B**) The proportion of CNV types (deletions or duplications) and (**C**) bases affected by CNVs for rhesus macaques compared to humans.

We find that macaque CNVs are distributed across all macaque chromosomes, but unevenly, with some stretches completely void of events and others where CNVs seem to be enriched (Figure 2). Contrary to previous studies in rhesus macaques (Lee et al. 2008), we find that the number of CNVs on a chromosome is strongly correlated with the length of the chromosome (Figure S1). This may again be the result of the increased resolution in our study. We also observe some clustering in the telomeric regions (Figure 2). This telomeric clustering is consistent with the duplication maps of the macaque genome (Gibbs et al. 2007) and the human genome (Bailey et al. 2001; Fortna et al. 2004; Zarrei et al. 2015), and is likely driven by the higher concentration of transposable elements in these regions, which mediates higher levels of non-allelic homologous recombination (i.e. unequal crossing-over).

**Figure 2:**
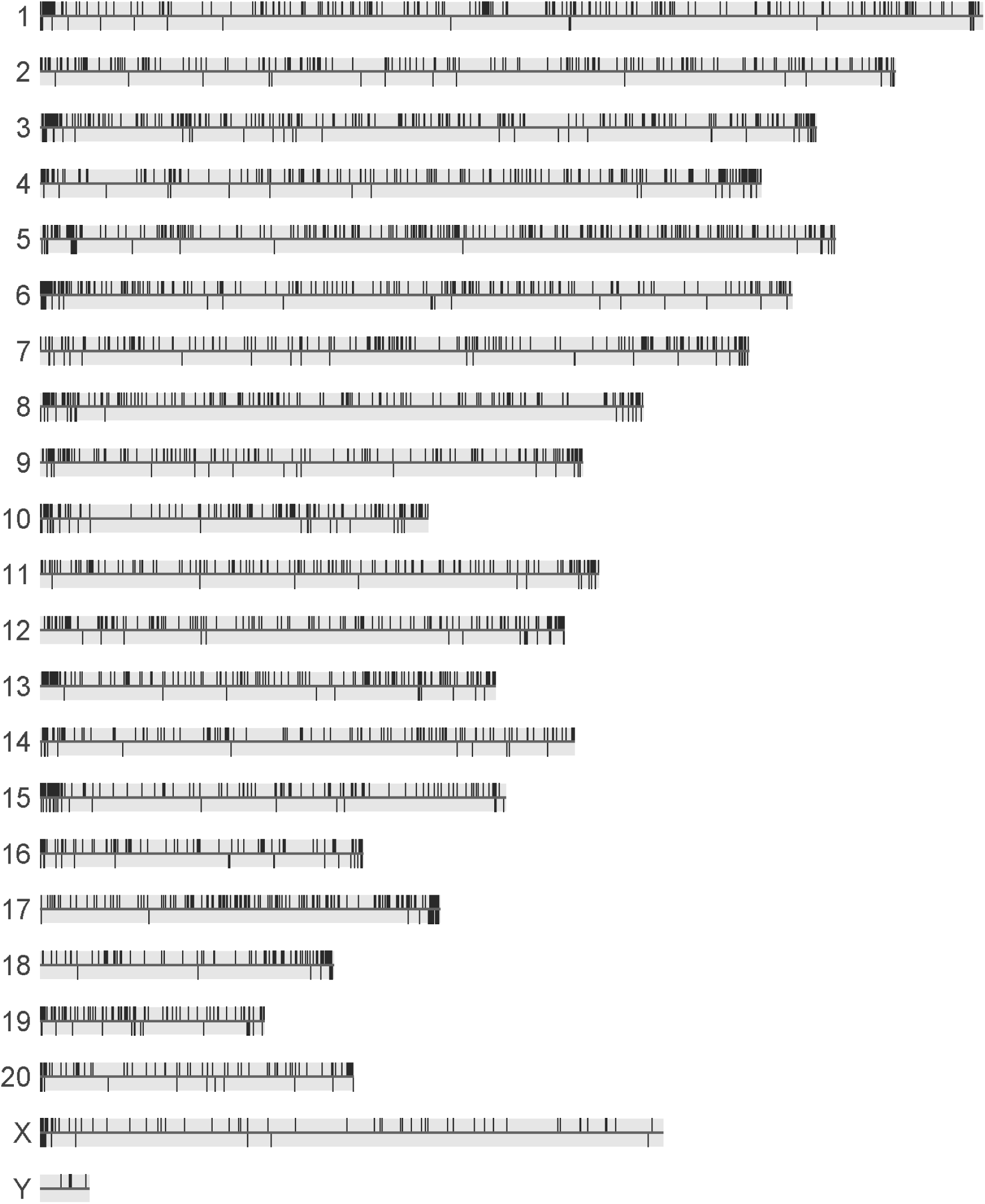
Locations of identified CNVs on the 22 rhesus macaque chromosomes. Deletions extend above the horizontal line for each chromosome and duplications extend below it.

We used published CNVs from a sample of 235 humans (Brandler et al. 2016) to study the similarities and differences between primate species. We find that the proportions of segregating deletions and duplications are not significantly different between the two species (Figure 1B; *χ*^2^ = 3.77, d.f. = 1, *p* > 0.05). Given the observed bias toward deletions, it is unsurprising that both species have a higher proportion of bases deleted than duplicated (Figure 1C). The average individual in our macaque sample is heterozygous for 1,317 CNVs that delete 2,804,650 base pairs (bp) and duplicate 471,100 bp.

CNVs in macaques have an average length of 3,519 bases, with duplications (mean length 6,853 bp; min length 138 bp; max length 97,301 bp) being longer than deletions (mean length 3,076 bp; min length 40 bp; max length 98,035 bp). Compared to humans, macaques have longer CNVs on average (Figure 3A; Kolmogorov-Smirnov *D* = 0.46, *p* << 0.01) and this pattern holds for both deletions (Figure 3B; Kolmogorov-Smirnov *D* = 0.48, *p* << 0.01) and duplications (Figure 3C; Kolmogorov-Smirnov *D* = 0.38, *p* << 0.01). It is unclear whether this shift in CNV length distributions between macaques and humans is a true biological phenomenon, which would point to some change in the underlying CNV mechanism, or simply reflects our inability to detect very small variants in macaques. We took every effort to eliminate methodological bias between the macaque CNV calls and the human CNV data set. In their paper, Brandler et al. (2016) use several different CNV calling and genotyping methods. We have restricted our comparisons to CNVs called with the same methods we employed for the macaque data, namely CNVs called with Lumpy (Layer et al. 2014) and genotyped with SVtyper (Chiang et al. 2015). To test the effects of different CNV calling methods and filtering steps, we made the same comparison between macaque and human CNV lengths while using the full human dataset (Figure S2) and without filtering the macaque CNV calls (Figure S3). Regardless of the partitioning method used, we still observe that macaques have, on average, longer CNVs than humans. Another possible technical explanation for this observation may be the sequencing libraries used in the two datasets: while Brandler et al. (2016) sequenced most samples with a read length of 100 bp and insert sizes of 113 bp, the read length of the macaque sequences was larger at 150 bp and an insert size of 128 bp. Although we would expect that this difference in read length would allow the macaque calls to be more sensitive to smaller events, it may play a role in the resulting length of CNV calls. It is also possible that the difference in length distributions is due to a still unidentified technical difference between the two studies.

**Figure 3:**
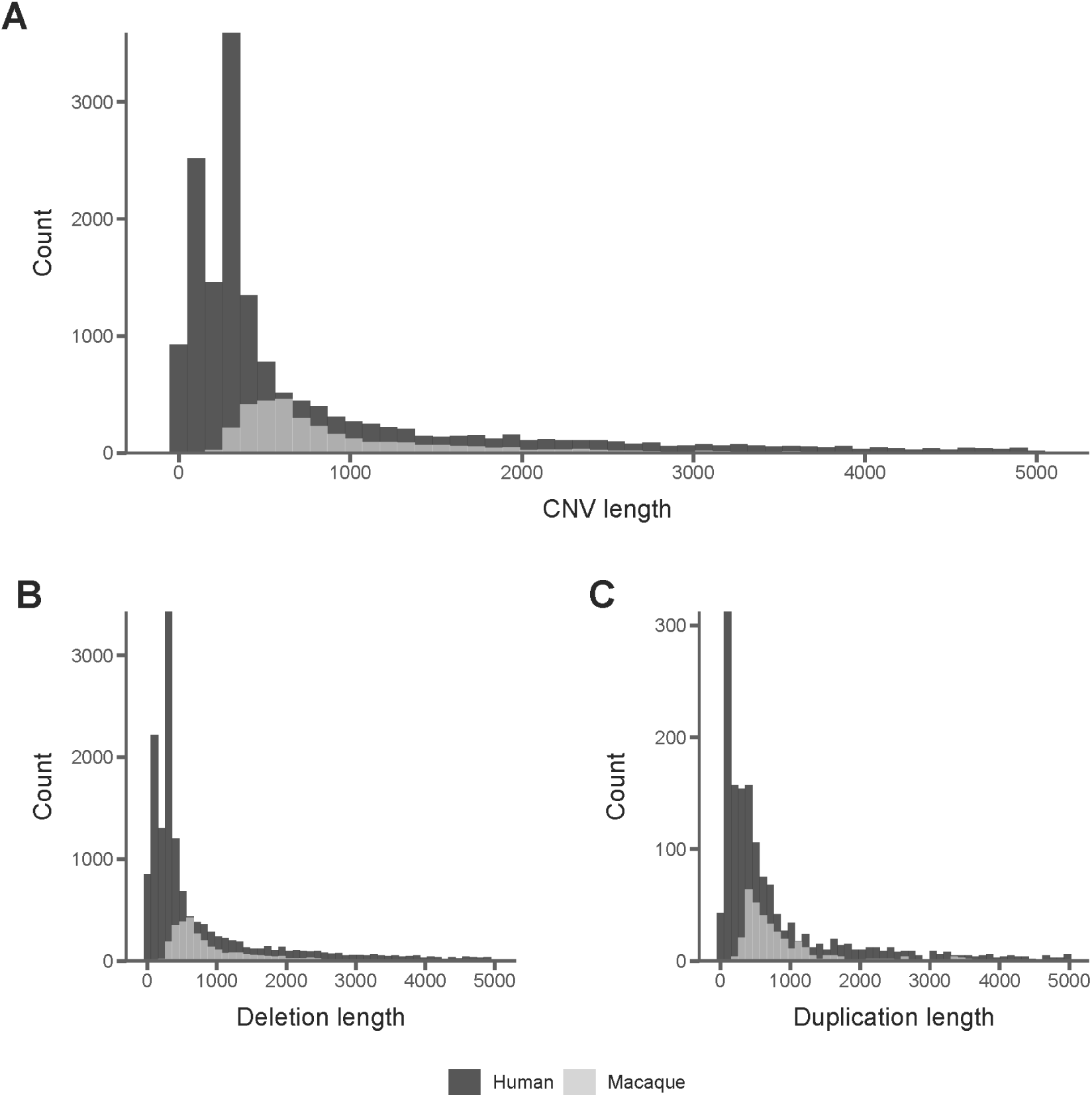
Length distributions of CNVs shorter than 5000 bases. Distributions shown for (**A**) all CNVs, (**B**) deletions only, and (**C**) duplications only.

### De novo *copy-number variants*

In a companion study we have described the rate and pattern of *de novo* single nucleotide variants in rhesus macaques (Wang et al. 2019). Here, we identify *de novo* CNVs in the same individuals by looking for CNVs that are unique to the offspring in a trio, as well as being in a heterozygous state. We find only 9 total *de novo* CNVs among our 14 macaque trios, consisting of 7 deletions and 2 duplications. This number of mutations makes the expected number of *de novo* CNVs 0.32 (95% CI 0.13-0.52) per generation per haploid genome. This rate of mutation is similar to that reported in humans (Brandler et al. 2016), which is consistent with the similar genome size between the two species. In contrast, the mutation rate of CNVs in *D. melanogaster* was found to be much lower (0.025 per genome; Schrider et al. 2013), though correcting for the ∼30-fold smaller size of the fly genome puts the mutation rates on the same order of magnitude per nucleotide.

By considering the age of sires when the offspring of each trio was conceived, we can ask whether the number of *de novo* CNVs increases in older fathers. We find no paternal age effect in macaques (Figure 4A; *R*^*2*^ = 0.033, d.f. = 12, *p* = 0.53), though with only nine events our study has low statistical power to detect an increase. However, we also performed the same analysis using 19 *de novo* CNVs from human trios (Brandler et al. 2016), and found no increase in the number of mutations in the offspring of older fathers (Figure 4B; *R*^*2*^ = 0.032, d.f. = 77, *p* = 0.12). Because the rate of new CNVs seems to be very low, increasing the sample size in macaques will increase confidence in our conclusion of a lack of paternal age effect in this species.

**Figure 4:**
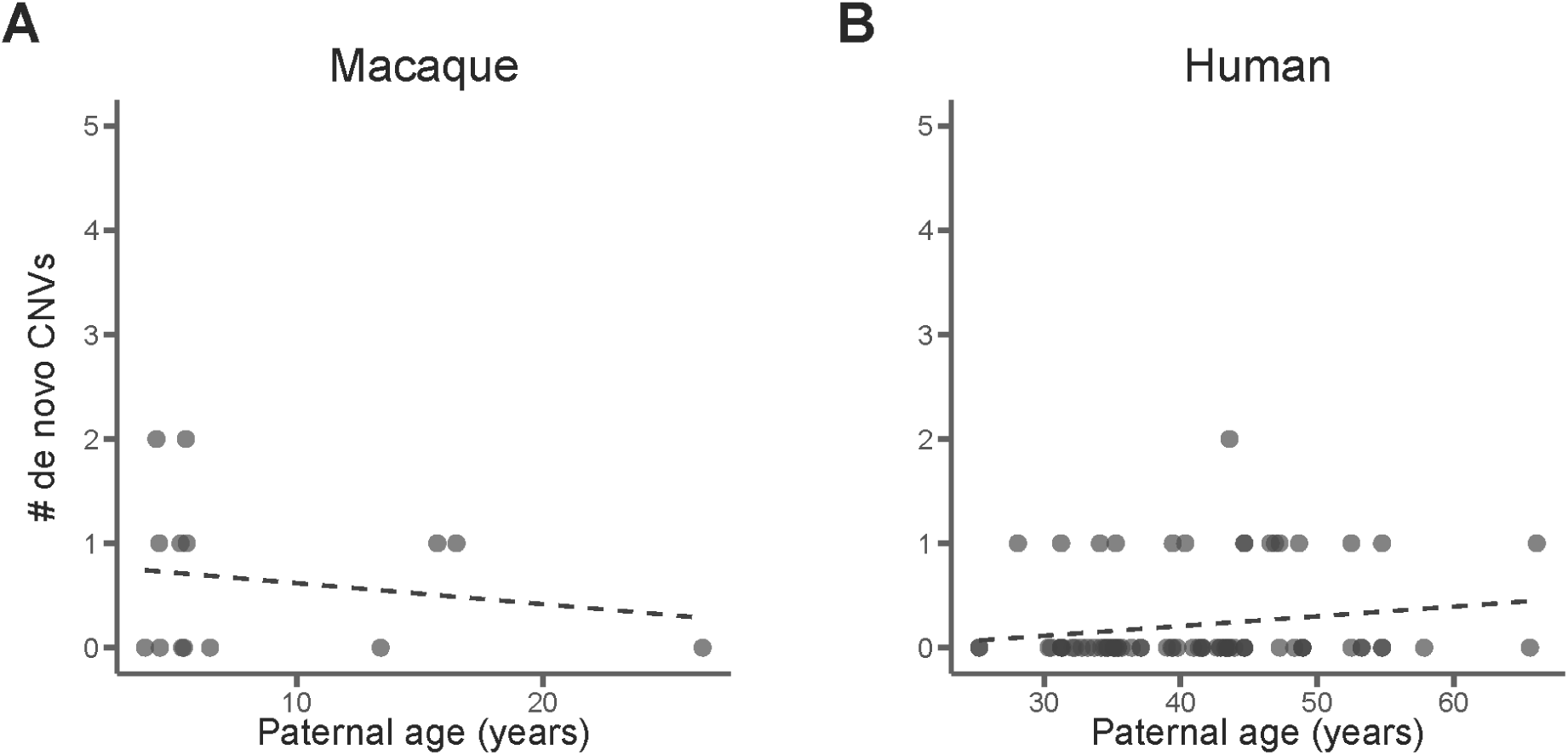
There is no correlation between *de novo* structural variants in (**A**)14 macaque trios or (**B**) 97 human trios (13 validated + 6 unvalidated CNVs). Each point represents a single trio.

### Gene duplications and losses within and between species

The ultimate fate of structural variants is to either become fixed in a population or to be lost. Genes overlapping CNVs can play a role in this process by conveying fitness benefits or costs depending on their copy-number. We investigated the long-term fate of genes involved in copy number variation in macaques using gene gains and losses among 17 mammal species (Figure 5A). By comparing the number of genes gained and lost between species to the number of genes overlapping segregating CNVs within macaques, we uncover patterns in short- and long-term evolution of gene copy number.

**Figure 5:**
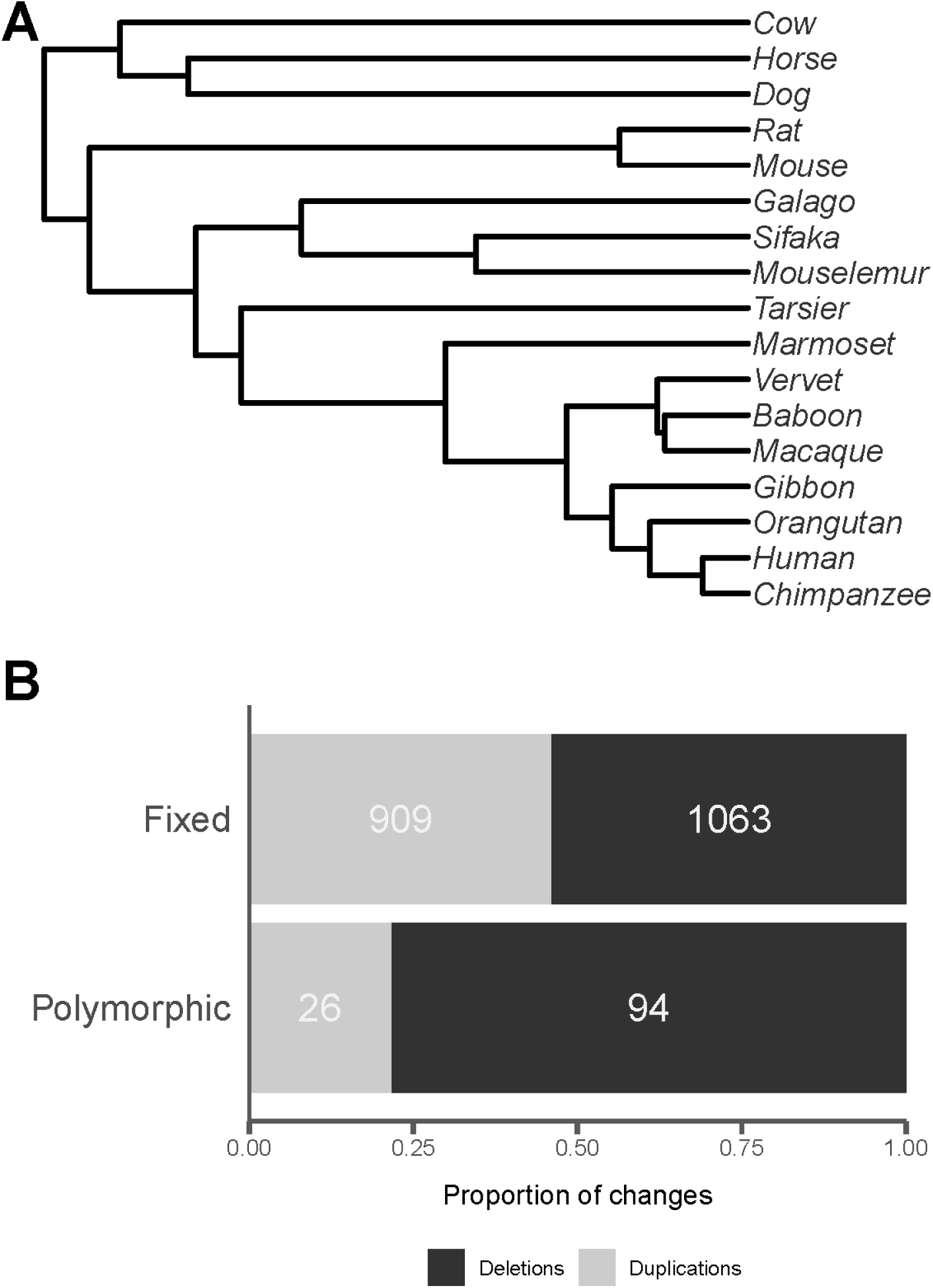
(**A**) Long-term patterns of gene gain and loss were inferred for macaques by comparing gene copy-numbers among 17 mammal species. (**B**) Among genes in both the gene family (Fixed) and CNV (Polymorphic) analyses, we find that genes are more likely to be part of polymorphic deletions, and conversely that there is a larger proportion of duplications among fixed differences.

We find that among the 3,754 CNVs in the macaque samples, 203 of them overlap 338 genes (out of 32,382 total annotated genes); the vast majority occur in intergenic regions. Of the CNVs that overlap a genic region, most span more than one gene (average 1.67 genes per event). However, this is driven by a few CNVs larger than 25 kb that overlap 2-3 genes. CNVs shorter than 25 kb overlap on average only 1.18 genes. This is the same pattern observed in humans, where, of CNVs overlapping genes, they include an average of 1.18 genes.

Among the genes within CNVs in macaques, 244 (72%) have been wholly or partially deleted, while 94 (28%) have been wholly or partially duplicated. The ratio of deleted to duplicated genes in macaques is 2.60, which is much lower than the overall ratio of deleted to duplicated regions (7.51). The over-representation of duplicated genic regions compared to non-genic regions has been observed previously in primates (Fortna et al. 2004; Dumas et al. 2007; Sudmant et al. 2013) and suggests that gene duplication is less costly in the short-term than deletion. The ratio of deleted-to-duplicated genes in macaques is also significantly higher than the ratio in humans of 1.46 (*χ*^2^ = 13.17, d.f. = 1, p << 0.01), possibly because of an increased rate of duplication in the great apes (Hahn et al. 2007; Marques-Bonet et al. 2009).

To examine the long-term fate of gene duplications and losses, we analyzed copy-number variation in 10,798 gene families across 17 species (Figure 5A). Along the branch leading to macaques since their common ancestor with baboons (∼11 million years ago), we infer the loss of 1,063 genes and the gain of 909 genes, for a loss-to-gain ratio of 1.17 (Figure 4B). This ratio is half that observed among segregating CNVs (see above) but could be biased because different genes may be included in the different annotation sets used. Restricting our CNV analysis to the 19,496 genes used in the gene family analysis, we find a ratio of 3.62 deletions to duplications, still significantly higher than the long-term ratio of gene gain to loss (Figure 5B; *χ*^2^ = 26.33, d.f. = 1, *p* << 0.01). Together, these results indicate that, while deletions dominate among *de novo* mutations and segregating CNVs in macaques, the number of genes gained and lost are more balanced over evolutionary timescales.

## Discussion

Copy-number variation can play a key role in disease and evolution (Eichler et al. 2007; Zhang et al. 2009; Girirajan et al. 2011). Here, we have shown that patterns of copy-number variation in rhesus macaques are largely similar to humans: segregating CNVs in both species are overwhelmingly made up of deletions. CNVs in macaques appear to be on average longer than in humans, though this may also be the result of an unidentified methodological bias. We found that *de novo* CNVs show no correlation with parental age in either species. This is in contrast to single nucleotide variants (SNVs), which have been found to increase with paternal age in both species (Kong et al. 2012; Jonsson et al. 2017; Wang et al. 2019). The difference between SNVs and CNVs is likely due to the differences in how these mutations arise. SNVs are thought to arise as errors in the DNA replication process during mitosis, or more rarely as unrepaired damage to DNA caused by the environment (Crow 2000). For male mammals, both of these processes are ongoing throughout the lifetime, with recurring mitoses occurring during spermatogenesis. However, copy-number variation is thought to arise only during unequal cross-over events during meiosis (Hastings et al. 2009; Zhang et al. 2009). Since meiosis occurs only once per generation, we expect no age effects for mutations that arise from it. This expectation is consistent with our present observations in macaques and previous studies in humans (MacArthur et al. 2014; Kloosterman et al. 2015).

With no age effect for copy-number mutations, we expect the rate of new CNVs per unit time (i.e. year) to be subject to a classic generation-time effect (Laird et al. 1969; Wu and Li 1985). The generation-time effect posits that species with shorter generation times accumulate more mutations over time because they experience more germline cell divisions per unit time.

This generation-time effect has been found to be dampened for single nucleotide mutations, which are dependent on mitosis, because of ongoing spermatogenesis (Thomas and Hahn 2014). However, for structural variants that occur during meiosis this effect should hold for neutral changes. The generation-time effect is a life-history model that provides a useful expectation when comparing rates of copy-number variation between species. Under this model, we would expect rhesus macaques, with shorter generation times, to have a higher rate of long-term copy number evolution than humans if life history traits are the only factor determining rates of copy-number changes.

Contrary to these expectations, the reverse relationship has been observed between species, with humans and chimps having the highest rate of gene gain and loss among primates (Hahn et al. 2007; Marques-Bonet et al. 2009). One possible explanation for the discrepancy between the expected and observed rate patterns of genic copy-number variation between these two species is a difference in selection between them. In this scenario, the underlying mutation rates per unit time differ, but studies of genic copy-number variation reveal the combined effects of mutation and selection in shaping the accumulation of change. In support of this is our observation in macaques that deletions make up the majority of polymorphic copy number events, but fixed gene gains and losses are more evenly balanced when comparing gene copy-number evolution between species. This is also further evidence for the claim that deletions are under stronger purifying selection than duplications (Conrad et al. 2006; Schrider and Hahn 2010; Schrider et al. 2013).

Taken together, the patterns of copy-number variation we have uncovered will help models of this type of mutation and their evolutionary consequences. While the patterns discovered here provide a good basis for this understanding, larger samples in future studies will provide higher confidence in quantitative estimates of parameters. Quantifying the rate of *de novo* CNV mutation may help us refine both disease models and the processes governing the evolution of the mutation rate. Being able to follow new variants from their introduction as mutations, to variation within populations, and finally to their fixation between species will reveal the evolutionary forces acting at every stage.

## Materials and Methods

### Sequencing and read mapping

Genomic DNA was isolated from blood samples of 32 rhesus macaques for whole genome sequencing (Illumina Nova-Seq, average 40X average coverage). Reads were mapped to the reference macaque genome (rheMac8.0.1, GenBank assembly accession GCA_000772875.3) using BWA-MEM version 0.7.12-r1039 (Li 2013) to generate a BAM file for each individual. Duplicate reads were identified with Picard MarkDuplicates version 1.105 (http://broadinstitute.github.io/picard/) and these reads were excluded from subsequent analyses. All BAM files were sorted and indexed with samtools version 1.9 (Li et al. 2009).

Reads that map to the reference with unexpected distances given their insert size (split reads) or orientations (discordant reads) between mate pairs can be used as signals of genomic deletion and duplication. These split and discordant reads were identified in each individual with samtools version 1.9 (-F 1294 for discordant reads) and the extractSplitReads_BwaMem script included in the Lumpy (Layer et al. 2014) software package. This resulted in three BAM files for each individual used as input for the CNV calling software listed below: all reads, discordant reads, and split reads.

### Calling copy-number variants (CNVs) in Rhesus macaques

Copy-number variants were called only on contigs that map to assembled macaque chromosomes. We used a suite of methods in the SpeedSeq software (Chiang et al. 2015) that use patterns of split and discordant read mappings to identify structural variant breakpoints throughout the genome to call CNVs. First, Lumpy (Layer et al. 2014) was used to find putative breakpoint sites in all 32 macaque individuals. Lumpy uses several pieces of evidence (such as split and discordant reads) to probabilistically model where breakpoints occur in the genome. CNVs called by Lumpy were genotyped with SVtyper (Chiang et al. 2015), which uses a Bayesian framework much like that used to genotype single nucleotide variants to determine whether CNVs are homozygous or heterozygous. For CNV calling with Lumpy, repetitive regions were masked using the rheMac8 RepeatMasker table from the UCSC table browser (Karolchik et al. 2004; http://genome.ucsc.edu/).

The software SVtools (Larson et al. 2018) was used to combine the calls from the 32 individuals into a single set. This set was then re-genotyped with SVtyper to obtain information for all CNVs in all samples (even if they were not present in that sample) for filtering. CNV calls were annotated with read depth information using Duphold (Pedersen and Quinlan 2019) and finally CNVs were pruned with SVtools such that, among events found to occur within 100 bp, only the event with the highest quality score was retained. CNVs were then annotated as to their overlap with genes by using the UCSC table browser. GNU Parallel (Tange 2011) was used throughout to parallelize the CNV calling software across individuals.

### Filtering putative macaque CNVs

The process for calling CNVs resulted in 157,914 events at 8,515 sites. To reduce the number of false positives, we applied the following filters to our set of CNVs:

1. Removed 83,371 CNVs at 2,615 sites that are present in at least 31 of the 32 individuals. These are most likely events in the reference individual, or misassemblies.
2. Removed 4,934 CNVs at 464 sites over 100,000 bp in length.
3. Removed 435 CNVs at 244 sites with a quality score less than 100.
4. Retained only deletions in which the fold-change of read depth for the variant is < 0.7 of the flanking regions. This filter removed 12,763 CNVs at 870 sites.
5. Retained only duplications in which the fold-change of read depth for the variant is > 1.3 of regions with similar GC content. This filter removed 9,954 CNVs at 568 sites.

These filters yield a reduced CNV call-set of 46,457 events at 3,754 sites which was used for all subsequent analyses.

### *Identifying* de novo *CNVs and calculating the mutation rate*

From the full set of 3,754 CNVs, we identified *de novo* events as those that occur only in one of the probands of the 14 trios. We required both parents to be homozygous for the reference allele and the child to be heterozygous. For F1 probands, the *de novo* CNV was allowed to be present in the proband’s offspring, as new mutations would be expected to be transmitted roughly half the time. This occurred in 2 out of the 3 F1 CNVs.

We calculated the CNV mutation rate per generation for a haploid genome by taking the mean number of transmissions in the 14 macaque trios and dividing by 2. Standard error for this rate was calculated by taking the standard deviation of the number of transmissions for the 14 trios divided by the square root of the number of trios times a critical value of 1.96 for the 95% confidence interval.

### Human CNV data

Human CNVs were downloaded from the supplemental material of Brandler et al. (2016). This study used 235 individuals in 69 families to look for patterns of *de novo* structural variation among autism patients. Their validated *de novo* mutations along with parental ages were obtained from their supplemental spreadsheet S1 and used for Figure 3B. The entire CNV call-set from their supplemental data S1 was used for all other comparisons. These authors used two methods to call CNVs, Lumpy (Layer et al. 2014) and ForestSV (Michaelson and Sebat 2012), and two methods to genotype their CNV calls, SVtyper (Chiang et al. 2015) and gtCNV (now known as SV^2^; Antaki et al. 2018). We restrict our comparisons to those called with Lumpy and genotyped with SVtyper for consistency with our methods.

### Counting fixed macaque gene duplications and losses

In order to identify genes gained and lost on the macaque lineage we obtained peptides from human, chimpanzee, orangutan, gibbon, macaque, vervet, baboon, marmoset, tarsier, mouse lemur, sifaka, galago, rat, mouse, dog, horse, and cow from ENSEMBL 95 (Zerbino et al. 2018). To ensure that each gene was counted only once, we used only the longest isoform of each protein in each species. We then performed an all-vs-all BLAST (Altschul et al. 1990) on these filtered sequences. The resulting e-values were used as the main clustering criterion for the MCL program to group peptides into gene families (Enright et al. 2002). This resulted in 15,662 clusters. We then removed all clusters only present in a single species, resulting in 10,798 gene families. We also obtained an ultrametric tree (Figure 4A) from a previous study (Rogers et al. 2019) for 12 mammal species and added mouse lemur (Larsen et al. 2017), tarsier, vervet, and galago based on their divergence times from timetree.org (Kumar et al. 2017).

With the gene family data and ultrametric phylogeny as input, we estimated gene gain and loss rates with CAFE v4.2 (Han et al. 2013) using a three-rate model, which has been shown to best fit mammalian data (Hahn et al. 2007; Marques-Bonet et al. 2009; Carbone et al. 2014). CAFE uses the estimated rates to infer ancestral gene counts and we subsequently counted the number of genes gained and lost in the macaque lineage relative to its common ancestor with baboon.

## Data availability

Genome sequences for the 32 rhesus macaque individuals are available on the sequence read archive (accession XX) (Wang et al. 2019). All other data, including pedigree structures, CNV calls, and scripts for reproducible figures and analyses is available as a GitHub repository: https://github.com/gwct/macaque-cnv-figs.

## Acknowledgements

We would like to thank Aaron Quinlan and Jonathan Beyleu for guidance in calling structural variants. We also acknowledge the members of The Mouse Lemur Genome Consortium for discussions of the mouse lemur genome used in the gene family analysis.

This work is funded by the Precision Health Initiative at Indiana University.

## Supplementary Figures

**Figure S1:**
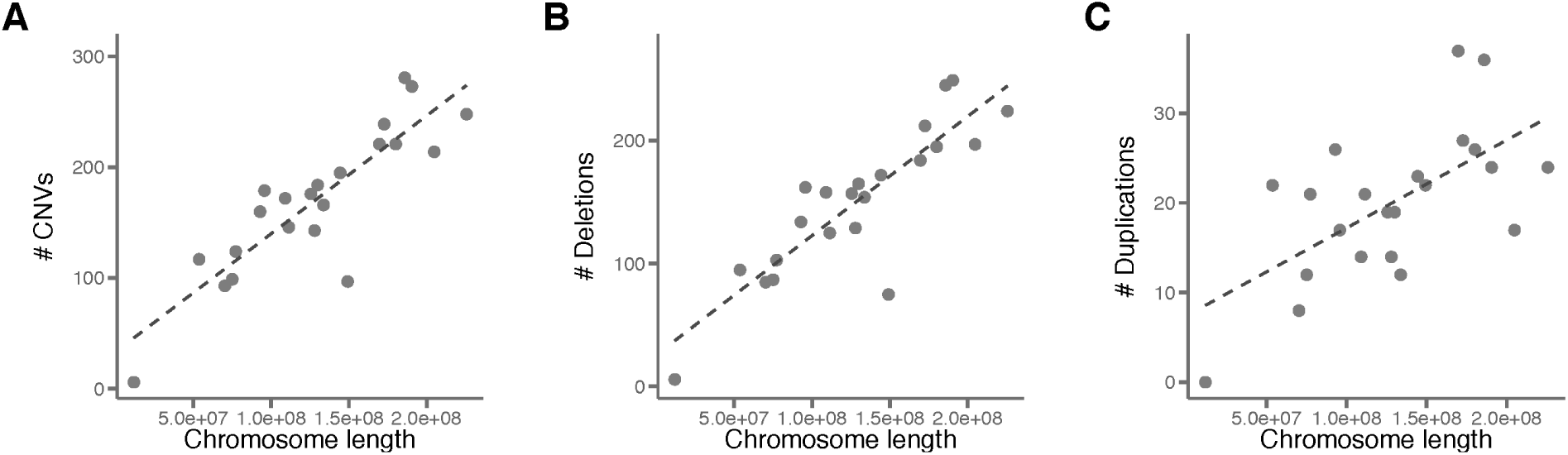
The number of CNVs is strongly correlated with chromosome length in macaques for **(A)** all CNVs, **(B)** deletions, **(C)** and duplications.

**Figure S2:**
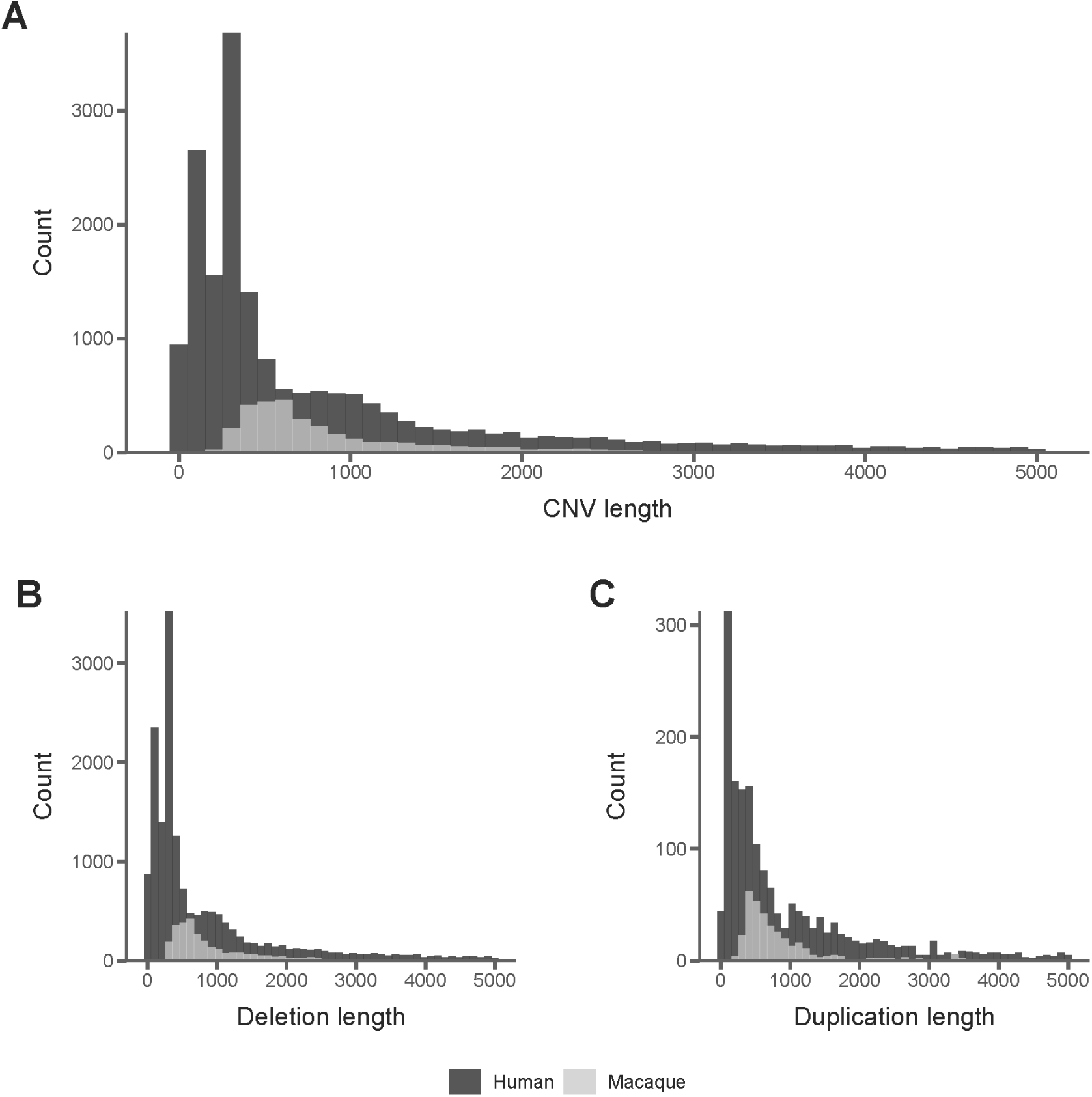
Length distributions of CNVs shorter than 5000 bases using the full human CNV dataset. Macaque CNVs are longer on average than humans for (**A**) all CNVs (Kolmogorov-Smirnov *D* = 0.43, *p* << 0.01), (**B**) deletions only (Kolmogorov-Smirnov *D* = 0.46, *p* << 0.01), and (**C**) duplications only (Kolmogorov-Smirnov *D* = 0.32, *p* << 0.01).

**Figure S3:**
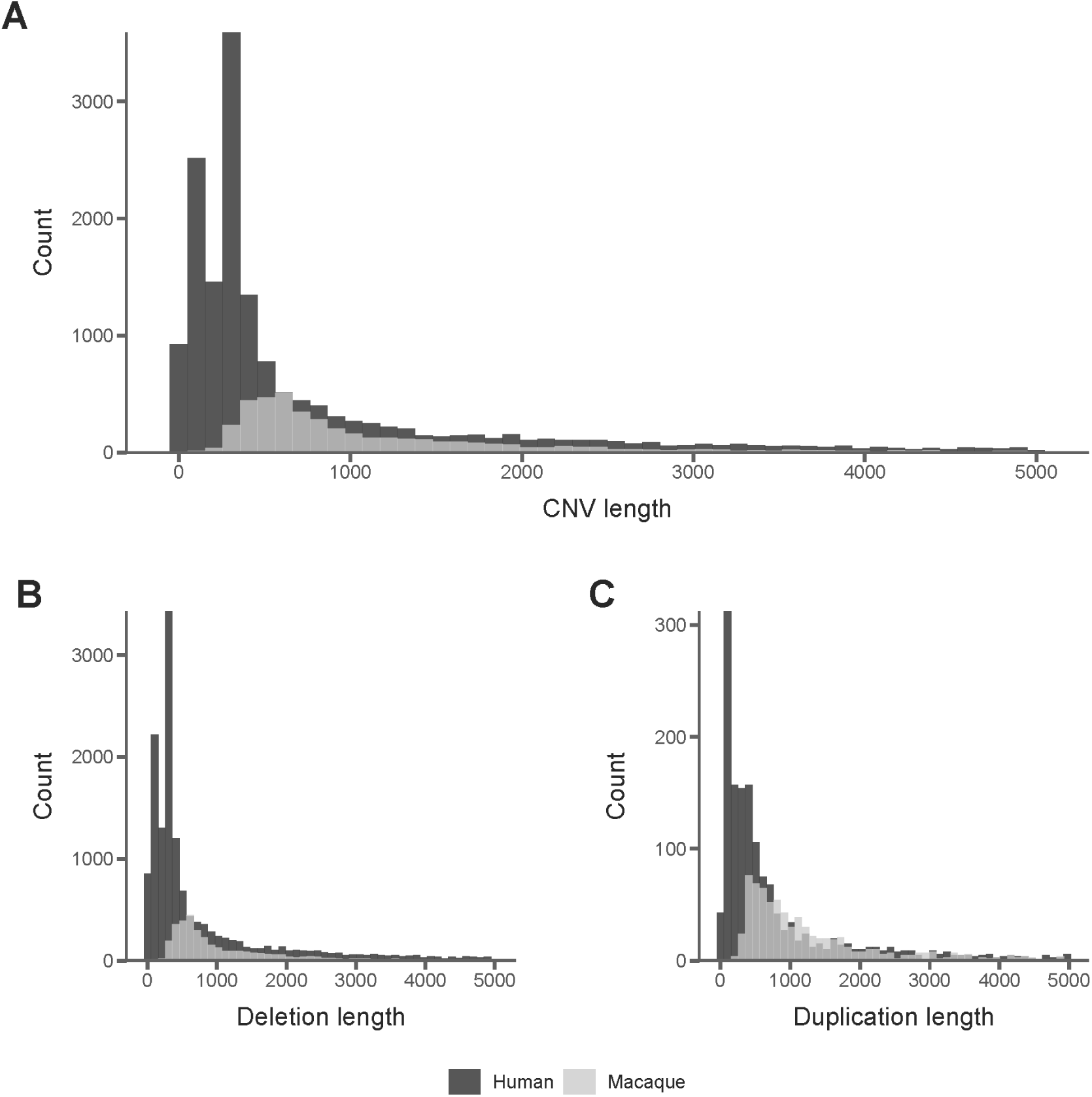
Length distributions of CNVs shorter than 5000 bases using the unfiltered macaque call set. Macaque CNVs are longer on average than humans for (**A**) all CNVs (Kolmogorov-Smirnov *D* = 0.47, *p* << 0.01), (**B**) deletions only (Kolmogorov-Smirnov *D* = 0.48, *p* << 0.01), and (**C**) duplications only (Kolmogorov-Smirnov *D* = 0.42, *p* << 0.01).

**Figure S4:**
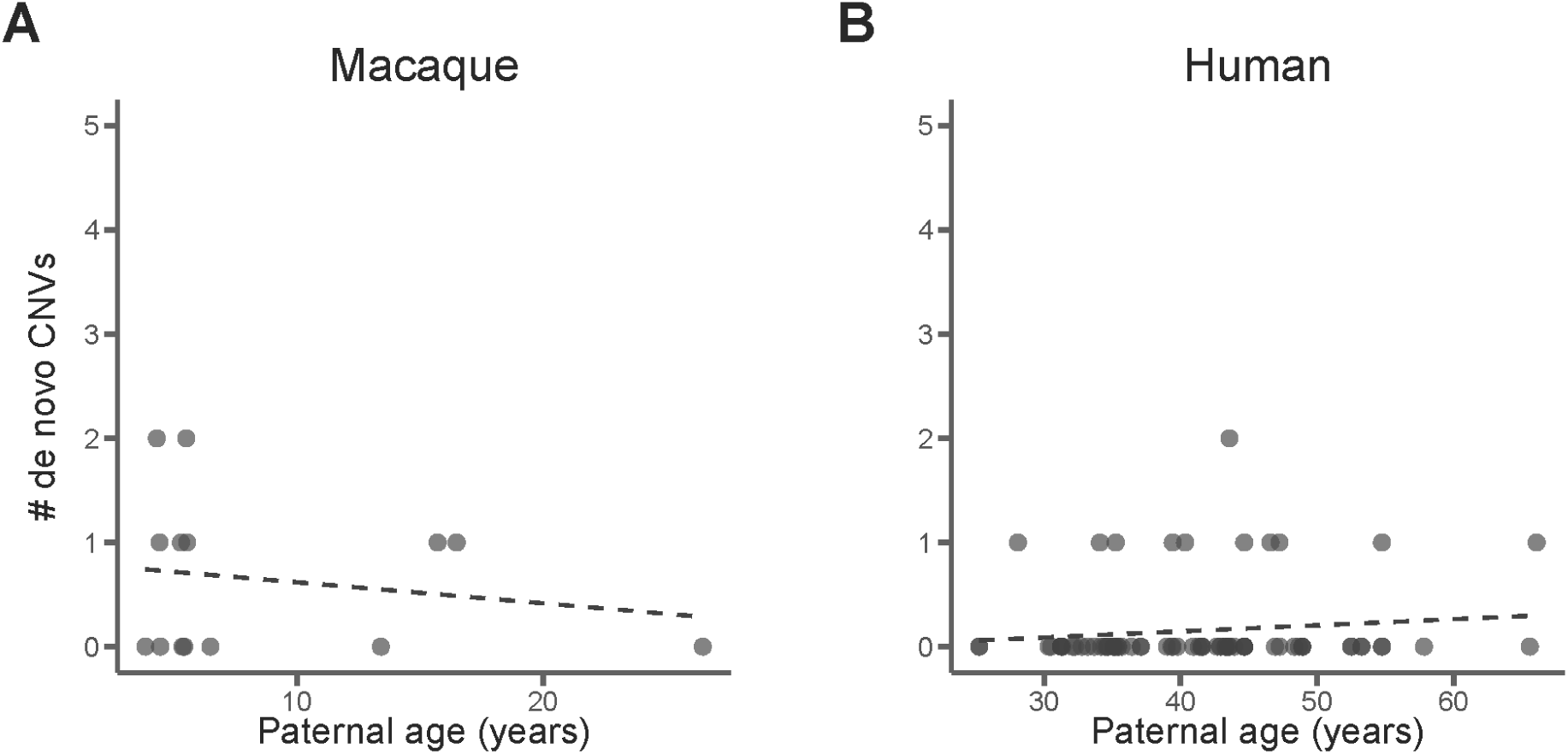
There is no correlation between *de novo* structural variants in (**A**) 14 macaque trios or **(B)** 97 human trios (13 validated CNVs only). Each point represents a single trio.

## Notes

https://github.com/gwct/macaque-cnv-figs

